# Structural and modelling insights into the dynamic association between the transcription factor and DNA

**DOI:** 10.1101/2023.10.13.562174

**Authors:** Feng Jin, Ke Xu

**Author notes:** Correspondences (K. X.).

## Abstract

The DNA recognition mechanisms by transcription factor (TF) was a significant scientific issue in the gene transcription and regulation. Multiple research technology including the experimental and modelling method has been introduced into the study of this aspect. In this article bioinformatic, protein modelling and dynamic simulation method was employed to display the overview of the dynamic binding between TF and DNA. Physical properties of positional change and freedom of atoms in addition with the volume exchange and the interaction analysis revealed the flexible binding sites of this element. The association of TF increased its stability with dynamic conformational change. The different levels of resistance to the sequential fluctuations of the residues and the nucleotides in the binding site stabilize the overall structure of the complex and initiated the open of the double helix that indicated the molecular mechanisms of the recognition and regulation of the elements.

## Introduction

The gene expression in various species is a highly regulated process that is involved with many different types of genetic elements and proteins which are associated with different types of DNA sequence. The proteins (usually named transcription factors) that are present in the DNA sequence recognization are typically classified into two types^1,2^. The first type of TF globally regulated most gene expression and recognize similar genetic codes^1,2^. The second type of TF specifically binding to the DNA sequence with distinct properties^1,2^. These TF usually contains DNA binding motifs with unique features, including zinc finger, HLH (helix loop helix), and so forth^3^. A typical zinc finger motif consists of two helix fingers and a zinc atom, which usually repress the gene expression in either prokaryotic and eukaryotic cells^3,4^. While a typical HLH motif was commonly constructed by two helix loops that linked by a short and flexible loop^3,4^. The investigation of DNA recognition mechanism dates back to centuries ago, yet remains a mysterious and unsolved scientific issue^1,2^. Various theories about how DNA matching evolved from many different scientific groups^1-3^. Many different methods are established and modern technology are available including DNA sequencing, genetic engineering techniques and other quantitative data collection tools such as SELEX ^2,5,6^. Structural analysis, statistical data and prediction tools provided large amounts of valuable information to facilitate the development of insights about the impact of the thermodynamic of the various molecules in the environment with or without the solution, including a particular type of elements on the TF recognization and stability analysis^2,7,8,9^. Besides, the following methods including protein modeling and dynamic simulation could be applied for display the entire or partial movement of the TF-DNA complex in femtosecond time scale and cover from picosecond to several microsecond^2,7,8,9^. However, the detailed molecular mechanisms of the different types of TFs from distinct families and the comparison of the identical and distinguish characteristics between them are not completely understood. Moreover, the initiation of the conformational change induced by the TFs and the principle underlying the behavior for the DNA recognition, associated motions and expectively the dissociation are still required further probing and investigation that makes the primary conscious of this research which are intended to be achieved mainly through the protein modeling and simulation method that improved from the conventional approach and concept. In this article, two TF families are selected given two categories that no structural information have been assigned so forth to determine their structural characteristics through protein modeling and provide their thermodynamic behaviors by dynamic simulation. Dynamic simulation method was firstly established by^7,8,10^ using Monte Carlo algorithm^7^ to create the first prototype of the new model system. Next, other methodologies^8^ were developed including Gromacs algorithms^7,10^ (also used in this article) and so forth. Following results were obtained, 1)The experimentally determined structures were analyzed to screen for potential example that covering the major families and possessing the typical properties, which is proper for the investigation of DNA recognition mechanisms and were used for the purpose of creating an dynamic behavior with dynamic simulation algorithm. 2) Two TFs with zinc finger motif without the assignment of entire structure were selected to predict the theoretical structure of the two TFs, DNA fragments they possibly recognized, and TF-DNA complex. 3) Dynamic simulation were performed through Gromacs algorithm by using the above experimental and predicted structures with the same parameters in 2fs time scale, running for 10ns and 50ns under constant temperature and pressure. 4) Parallel simulation with the same parameters and conditions as in the previous simulation with the identical model and algorithm were performed, achieving the same results. 5) The simulation results were analyzed to identify the dynamic properties of the above three elements including the movements of the structure and stability analysis that were represented by the positional change, flexibility reflected by the statistical freedom of atoms per residue in proteins or base in nucleotide of the DNA fragments, solvent accessibility surface area and the binding properties of the TFs and DNA fragments. In conclusion, the results of these analysis suggest the different impacts of the different types of TFs on the stability of TF-complex and dynamic binding between TF and DNA, moreover, the dynamic interaction between these TFs and DNA codes were discovered to display the characteristics of these elements as well as their ability to distinguish between different types of DNA sequences from other types of genetic elements.

## Results

### The DNA association of the transcription factor Rex stabilized the structure of the TF but initialized the conformational change and the open of the double helix strand

The first step in the development of the dynamic overview of the transcription regulation is to establish the framework of the implementation and the structural principles for the plenipotentiary of the entire system. It is in the context of the gene transcription process in the field of the molecular simulation and the dynamic inspection into the DNA structural regulation and the binding mechanism for the various types of the transcription factors in the genome of prokaryotic and Eukaryotic species. These include the deposits of the experimentally determined crystal structures and the predicted models into the dynamic simulation process that performed under the suspect simulation system of particular conditions described in the method section and listed in the supplementary table S1.

This article employs the developed Gromacs 2022.1 program to perform the simulation process under the constant temperature and pressure conditions. It imitate the experimental condition in the laboratory environment and provide the necessary algorithms, numerical equations as well as functional information for the generation of the statistical movements of the atoms in the production of the simulation results. The results were displayed in the format of line plot and structure overlap between the different simulation points on the Fig. 1-4 and the supplemental figure S1 as named Fig. S1. The configuration was saved every 10 ps with the overlapped configurations from different snapshots output in 5ns time interval displayed by the specialized software mentioned above was shown in Fig. S1. The overall results were summarized in the following paragraph. It was revealed that the structure of the protein was stabilized by the association of the DNA reflecting from the fluctuation of the RMSD and so forth during the entire simulation process, indicating the structural movement between the initial and final configuration during the simulation process in 10ns and 50ns respectively.

**Figure 1.**
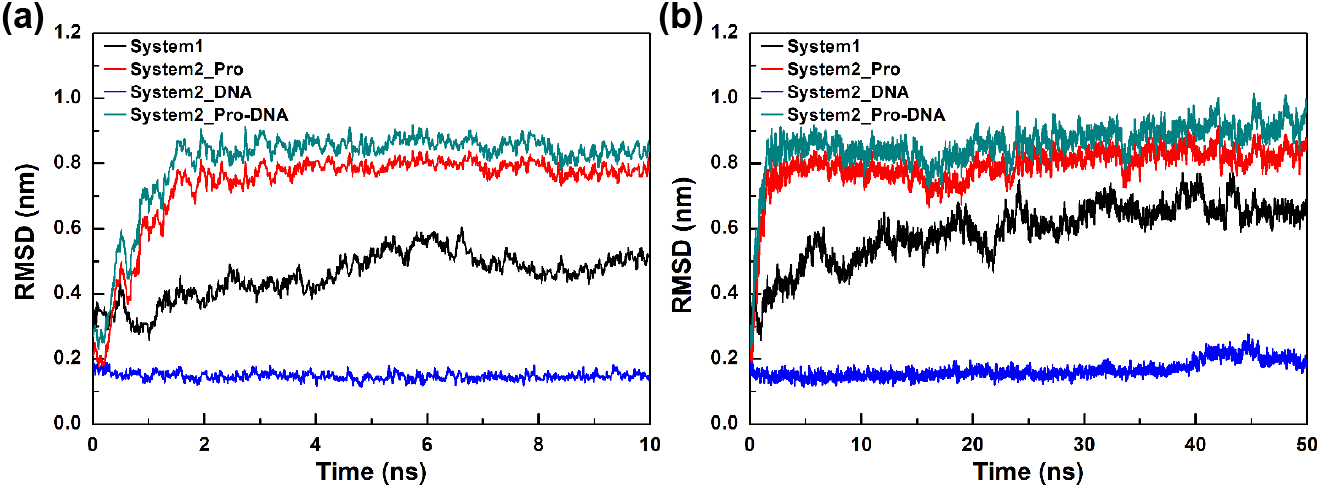
The RMSD of the protein alone, protein, DNA and the DNA associated protein complex in corresponding with the simulation time of 10ns and 50ns intervals. (a) The RMSD fluctuations of the protein only system and the protein, DNA and DNA associated protein complex vs the simulation time were shown in black, red, green and blue respectively in the 10ns time intervals. (b) The simulation time of the above systems in 50ns time intervals were shown in the same colors as in a. The protein alone system means in which the protein is not associated with DNA

The main steps of the project which is to create the first phenotype of the transcription factors behaved in the transcription regulation of the Eukaryotic genome was incorporated into the simulation process as follows. The initial state obtained from the experimentally identified crystal structures or the theoretical models were assigned to energy minimization by the steepest descent method of the installed package or the programs developed by the collaborate scientists to minimize the potential energy of the systems. The energy minimized state generated by the initial step was then transferred into the next stage of thermal equilibrium achieved by heating and cooling of the system in the atmosphere scale by the electromagnetic field and the energy optimized functions in this process with the efficient energy loss of the system as to approach to the thermal stable state of the system. This was fundamentally a prior process of the transition into the thermodynamic simulation of the universally organized system in the optimal conditions. It included the target protein or DNA as well as the potential magnitude volume of the solvent molecules additional with the ions that maintain the ionic properties of the liquid molecules producing the energy fluctuations in correlation with the atomsphere quantum in the numerically calculated strength for the energy force field of the system. This was further explained by the fact that the energy supply of the system was not sufficient for the purpose of generating electricity and thermothresholds of the system in the first circle and requiring more equilibration time circles to operate in parallel for the generation of the energy optimized state of the system for the entire time interval of 1ns. The resulting output of the system was much more appropriate for the purpose of the energetic equilibration of the system than the original state. The statistically predictable of the thermodynamic effects of the system was speculated by the Gromas package that stimulated the transient state of the system in the Gas function by increasing the energy density of the atoms that were generated by the electron beam and the energy functions. These properties were then applied to the electron magnitude changeable assignment to determine the maximum angle and distance exchange between atoms in the electron spectrum of the magnetic field to determine the radius distance of a photon quantum particle in the magnetic in charged systems. This was dynamically transmitted into the configurational change of entire atoms in the system that was able to be reflected to the conformational exchange in the thermal fluctuation circumstance that was detectable statistically by the specialized softwares and output in the plasma of the atoms in different format of chart for the representation of the quantum reflaction of the different types of molecules interpreted in the structure of the protein, DNA as well as the other molecules surrounding the solutes. The first properties of the simulation results were that of the conformational change of the protein which was evaluated by the positional dynamics of the entire molecule. It was in the statement of the root mean square deviation (RMSD in short) that describes the position of each atom in the molecule statically in the overall shape exchange of the molecules in the quantum speculative systems as shown in Fig. 1. The RMSD fluctuations of the 10ns and 50ns interval of the simulation time were separately recorded in a and b chart of this figure in which the average value of the RMSD was speculated from the current level of the data set that were collected from the previous described simulation process with the relative frequency of the fluctuation in the representation of different colors that were black, red, blue and green respectively in the response of the components in the protein only or DNA fragments included systems as indicated in the figure. The average RMSD fluctuations were 0.593±0.089 nm for the first system of the TF, Rex from *Streptococcus agalactiae*, or group B streptococcus (GBS), were less than that for the DNA associated system which was 0.789±0.73 nm, indicating that the stability of the protein was affected by the protein DNA interactions. This was due to the potential resistance of the protein to the thermal fluctuations of the system induced by the quantum inflation upon the nucleotides matching with the potentially volatile residues of the TF that was literally not quite tremendous for the thermo quantitative infections of the atoms in the systems. This was further evidence for the existence of the nucleotides in the universally recognized form of the protein-DNA complex for the transcription factors to be of the much more complicated and sophisticated nature than the original physically unstable system that included the protein only. Since the protein only system was initially unstable and fluctuated in the increasing conformational change reflected by the constant expansion of the protein in privilege by the value of the RMSD. However, the protein-DNA associated system is initially far more fragile than the protein only system but rapidly approaching to the stable state of the highest concentration of the protein conformational change within the first 5ns time interval and consequently underlying sufficient resistance to the thermal impact that solely resulted in frequently transmission of the protein configuration surrounded by the fast decay of the protein instant form to replicate the protein instance of the cell in the transcription initiation system that reflected by the surrounding RMSD within that of the average value.

The magnitude spectrum of each atom in the molecules that are present in the target systems were evaluated by the physical properties of root mean square fluctuations in short named of RMSF (Fig. 2) which were speculated by the atomic freedoms of each residues (a and b) in the protein or the nucleotides (c and d) in the DNA fragments of the protein-DNA complex. The fluctuation of the atoms were presented by the specialized softwares mentioned above which were reflected by the magnetic properties of the overall movements of the atoms in a particular amino acid residues or the base pairs in the nucleotides of the DNA fragments that were presented by the atmosphere fluctuations correlated with the amino acid residues or the base pairs of nucleotides in the DNA sequences. The nucleotides are normally considered to be less flexible as constantly contained in the double helix. Upon the initiation of the TF binding the freedoms of the atoms in the amino acid residues is further constrained by friction of the pairing bases in the interacting nucleotides that contributed to the atmosphere relationship between the nucleotides and the amino acid residues. The nucleotides of the DNA fragments exhibited slightly increase of the ionic induced flexibility in the unassociated forms as commonly considered in the former research (data not shown). However, the arise of atomic freedom for the bases of the nucleotides initiated by the surrounding atoms from the interacting amino acid residues and neighboring atoms in both the residues and nucleotides, as well as the induced conformational changes by the compacted atoms in the TF. Especially, the overall statistical freedom increase of the bases were in large correlations with the concentration of the population of the atoms that were condensed in the interacting amino acid residues. It is interesting to be pointed out that the density of the atoms in the largest freedom was condensed in the most flexible regions that in the linkage between the two helix concentrated functional domains that were involved in the DNA recognition and NAD^+^ binding, responsible for the implementation of the conformal change of the TF particularly upon DNA association. The less freedom of the atoms in this linkage in the 10ns time interval indicated the less far apart from the stable state of the initial experimentally determinate structure but a rational conformational change due to the flexibility increasable of the linkage induced by the additional adherence of the DNA fragments. This was partially metaled by the dimerization or complex formation within other transcription regulators in probably. The main difference between the two systems was that the electron density of the molecule was reduced due to the lacking of DNA. The protein alone system was colored in black while the protein-DNA complex system that evaluated the two components of the protein and DNA as well as the integrated one were shown in colors as red, blue and green respectively.

**Figure 2.**
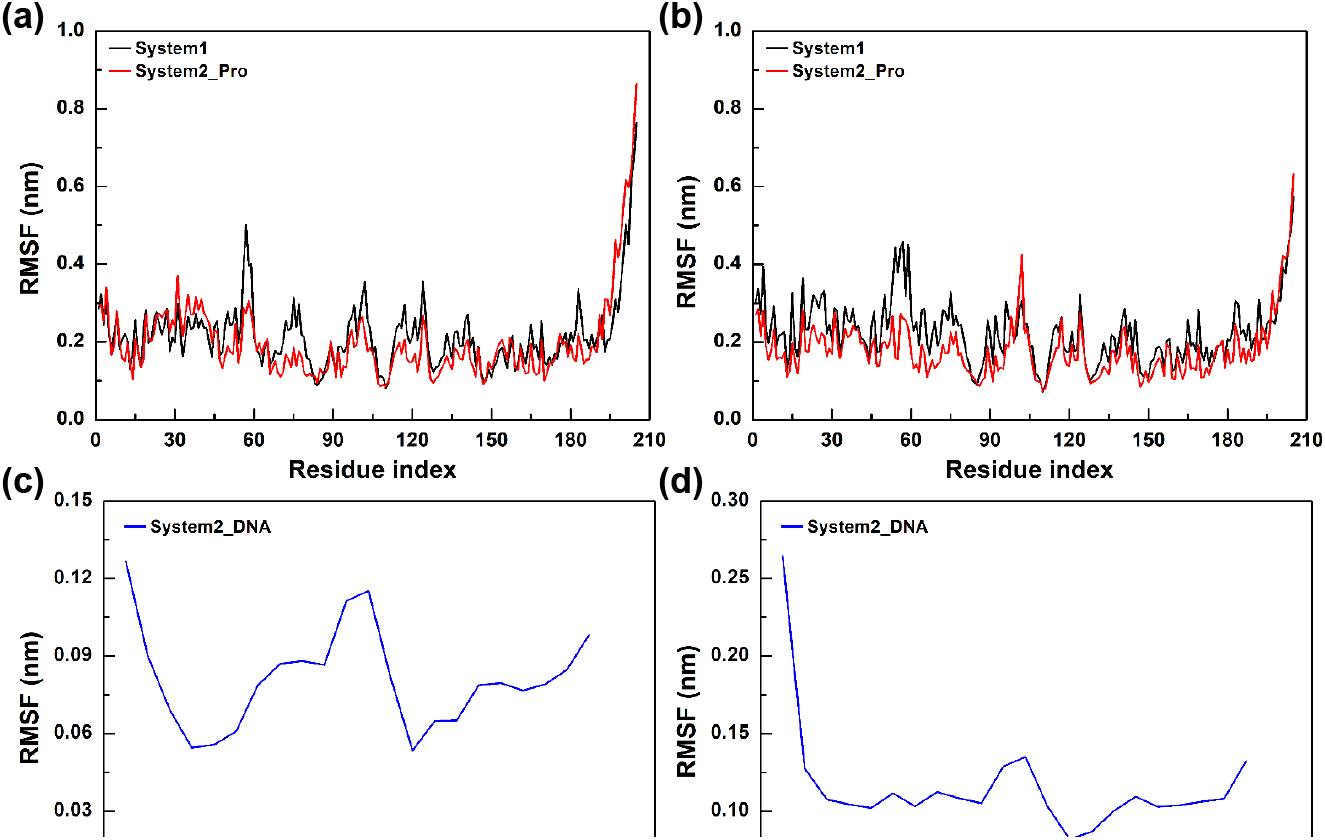
The RMSF distribution of the protein alone and the protein-DNA complex system in the time intervals of 10ns and 50ns respectively during the dynamic simulation process. (a,b) The RMSF distribution curve of the amino acid residues in the protein only system vs the simulation time was shown in black while that of the protein-DNA complex system was shown in red curves, in the time intervals of 10ns and 50ns respectively. (c,d) The simulation results of the bases of each nucleotides in the DNA fragments in the form of RMSF distribution in the 10ns (c) or 50ns (d) time intervals are shown in blue curves.

### The DNA association with the transcription factor Rex stabilized the initial structure provided by the volume exchange information on the structural transformation during the simulation process reflected through the solvent accessibility surface area (SASA) of the protein or the DNA bound protein

To further investigate the effects of the DNA association, the other properties that characterized the genetic regulation by the transcription factor Rex, such as the solvent accessibility surface area were immobilized by the identical software referred to in the first paragraph. As shown in Fig. 3, these properties were given in visualization by the line plots in different colors that represented the protein only (black) and protein-DNA associated systems including the protein (red), DNA (blue) and the complex (green). The DNA association did bot severely changed the volume and the fluctuation of the volume indicating that the DNA recognition process did not undergo severe volume change of the entire molecule. Consequently the average fluctuation of SASA for the protein is 119.885±4.427 nm^2^, which is smaller than that for the protein-DNA associated complex as was 120.613±3.819 nm^2^, indicating that the DNA recognition process also surrounded by the stability effect of the regulation elements in the genome reflected from the DNA fragments experimentally identified or predicted as the mimic of the DNA binding site, partially in consistent with the results of RMSD on the stability of the complex upon DNA binding.

**Figure 3.**
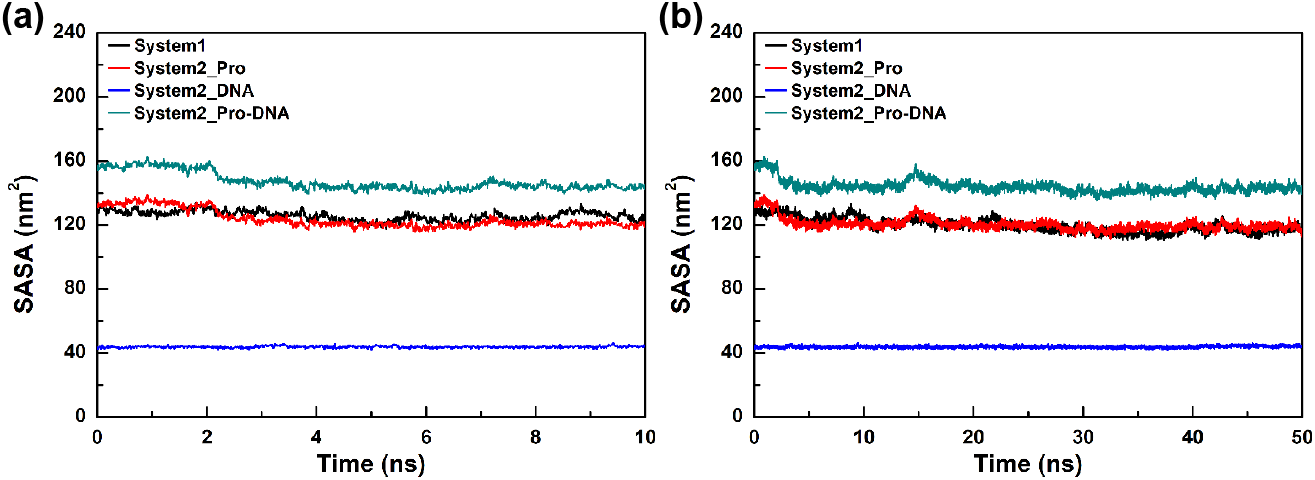
Shown is the solvent accessibility surface area for the protein without the DNA alone or protein-DNA complex in the 10ns or 50ns time intervals during the simulation process. (a) The SASA of the protein alone system is show in black while that of the two components and the protein-DNA complex were shown in red (for the protein) and blue (for the DNA) and green respectively during the simulation process with the time interval of 10ns. (b) The SASA of the above systems were shown in the same color as in a but with different simulation time interval that is 50ns.

### The transcription factor ZFH1 and ZFH2 was constructed with Alphafold to assign its theoretical structure and to predict its binding site respectively with the online software and database consequently evaluated with the dynamic simulation method exhibiting distinct properties compared with that of the Rex regulator

The dimerization of each interaction pair composed of amino acids and the nucleotides of the two components involved in the formation of the protein-DNA complex in the DNA binding site of the regulation elements which are the most commonly identified elements in the Prokaryotic and Eukaryotic cells were analyzed individually from the dynamic simulation data sets of 50ns time interval as shown in Fig. 1-4, with the most stable one shown in Fig. 4. The conformational change exhibit the DNA association and open process. Among these amino acid residues the LysX (Lys9, e.g.) residue behaved more flexible and dynamic than the others, since the amino acid residues approached to the 5’-terminal block the structural movements, this has been mentioned by the author that originally published the structure. Moreover the protein residues that directly located in the major groove of DNA also exhibited large conformational change and some are transiently preserved in the simulation time point passed through the equilibration stage in the simulation interval of 50ns.

**Figure 4.**
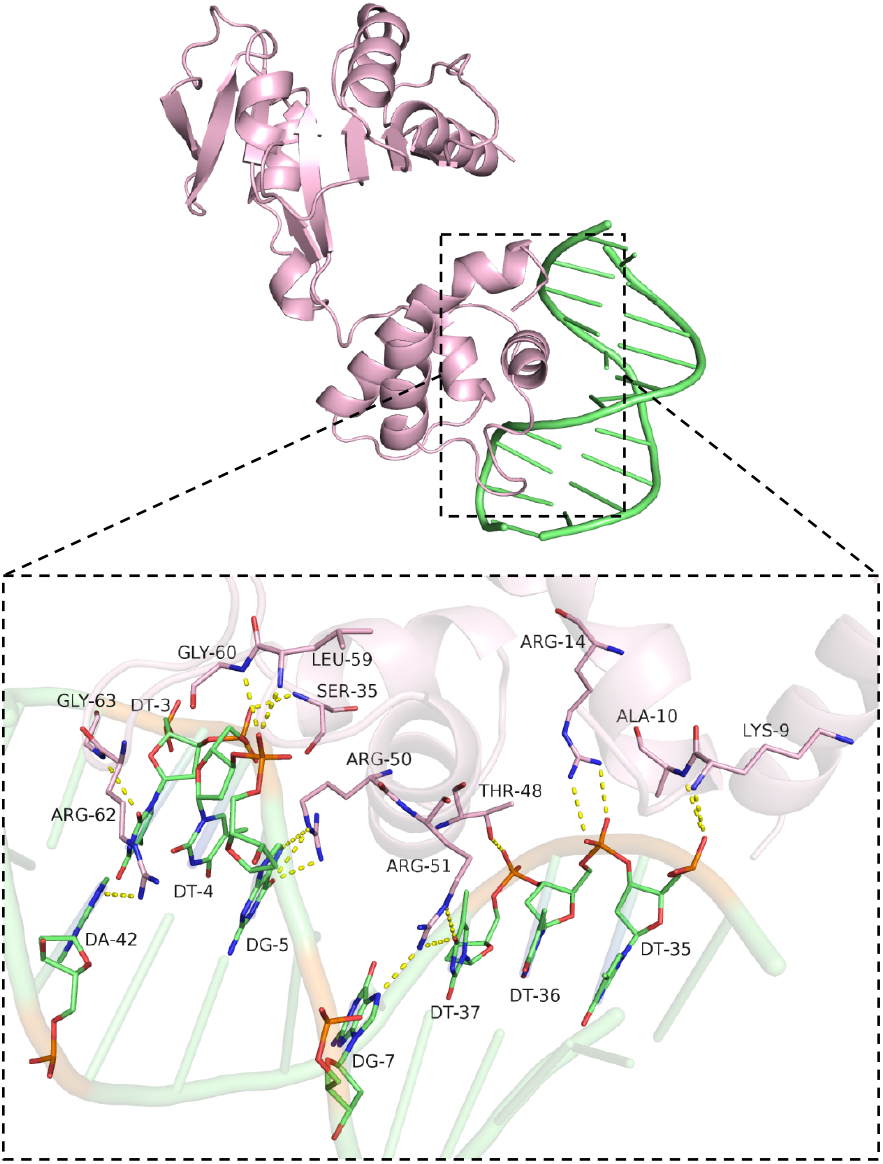
The structure of the transcription factor Rex associated with the corresponding DNA fragments, focusing on the individual amino acid residues and nucleotides interacting pair in the binding site. The structure of the DNA associated protein complex in cartoon form in which the protein residues and the DNA base pairs that contributed to the formation of the complex most but with highly dynamic were marked and shown in stick form with the bond displayed in dashed line between the individual interaction pairs of the protein and DNA fragments. Shown is the most stable conformation of the complex during the entire simulation process in which some interaction pairs were not observed in the initial stage.

In the case study of the other type of the transcription factors, two of them belonging to the C2H2 zinc finger family were selected and assigned for the protein modeling to construct the predicted structures since they were not determined experimentally. The researchers designed the research protocols to firstly assess the impact of the thermodynamic on the structural stability of the above TFs and then predict the potential DNA binding site in regarding with the existed experimental large amount of data set based prediction method provided on the http://zf.princeton.edu/form.php website that was provided by the research group from the university in preparation of the possible DNA fragments for the establishment of the theoretical structure of the DNA associated TF complexes through the Docking method including but not limited to the ZDock algorithm developed by the Scientists from the company mentioned in the contribution section. The above modeling results including the models for one of the two transcription factors, (Fig. S1), while the predicted DNA sequences that simulated the regulation elements that probably recognized and bound by the two TFs were listed below in the supplementary table (Table S2). As shown in the above figures, the TFs were composed of multiple functional domains that were linked by variable long loops which were identified to be of less similarity with the other types of the transcription factors. The DNA sequences potentially located within the DNA binding site were in large of similarities with each other but the key interacting pairs were not identical, indicating they were potentially regulating different genes that with slightly different regulating elements although they were belonged to the same transcription families with slightly differences particularly in the DNA binding regions. The two transcription factors were associated with the DNA major groove with the zinc finger domains that preferentially differentiate from the other types of the transcription factors with the similar DNA binding motifs. The DNA association mechanisms were discussed in the research articles published before but without structural analysis and undistinged DNA binding site leading to the formulation of the investigation in this article through the prediction and dynamic simulation methods.

The theoretical structure of the above two transcription factors were added to the simulation process that using the identical parameter and algorithms with the previous one described in the first section. The properties such as RMSD (Fig. 5), RMSF (Fig. 6) and SASA (Fig. 7) were also evaluated in similar format as that shown in Fig. 1-3. The RMSD of ZFH1 displayed in Fig. 5 was not quite fluctuated like those in Fig. 1 no matter for the 10ns or 50ns interval of the simulation time as well as that of ZFH2. Similarities of these two TFs belonged to the same family were foreseen in the prolonged simulation period according to the existed trend of the two proteins. However, different fluctuations of the RMSF for these two TFs were prohibited not only in the 10ns but also in the 50ns time interval as that were shown in Fig. 6. The ZFH1 were more flexible than that of the ZFH2 in most of the region and particularly in some of the domains including the DNA binding region that contain the C2H2 zinc finger motif while the ZFH2 were less flexible entirely but extremely freedom in the random loop included region. More interestingly, the solvent accessibility surface area of these two proteins were continuously decreased as displayed in Fig. 7 in both the 10ns and 50ns interval of simulation time distinguish with that for the Rex regulator. Regarding to the RMSD fluctuations in Fig. 5, this volume change of molecules were not due to the less stable state of the initial stage predicted but probably because of the large amount of the flexible coils that were compacted by the bulk molecules consist of water and ionics as described in the method section. The above results indicated that the stability of the two proteins are increased in the first 15ns and gradually approach to the steady state with less frequency of fluctuation. The average values of RMSD for ZFH1 (2.575±0.391 nm) was less than that for ZFH2 (2.788±0.409 nm), reflecting more stability of ZFH1 than ZFH1 although they were belongs two the same family and with relatively equal sequence identity, that potentially interpreted detailed functional dysclassification. It is consistence with the distinct RMSF distribution of the two TFs suggesting different overall flexibility but similar atomic freedom in the DNA binding domain, suggesting different functional characteristics that involved in the recognization of different elements with slightly sequence similarity but identical motif of highly conservation.

**Figure 5.**
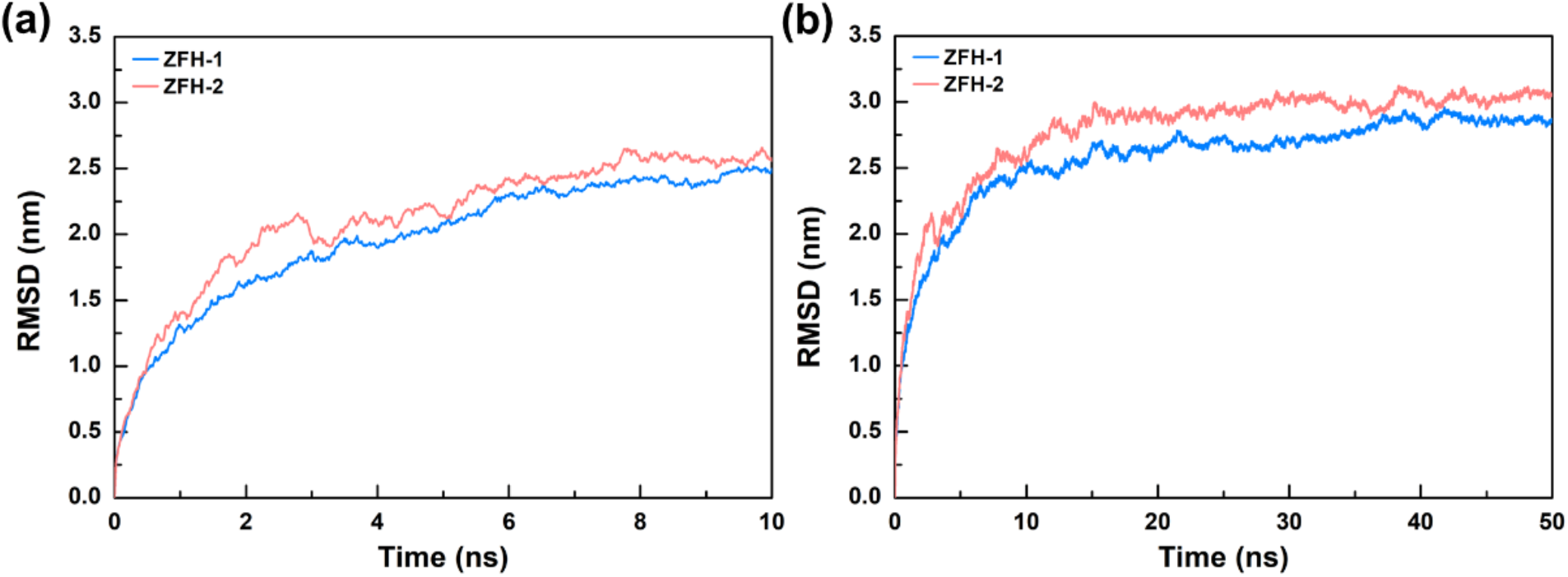
The transduction of the transcription factors which interpreted by the curves in different colors for ZFH1 in blue and ZFH2 in red.

**Figure 6.**
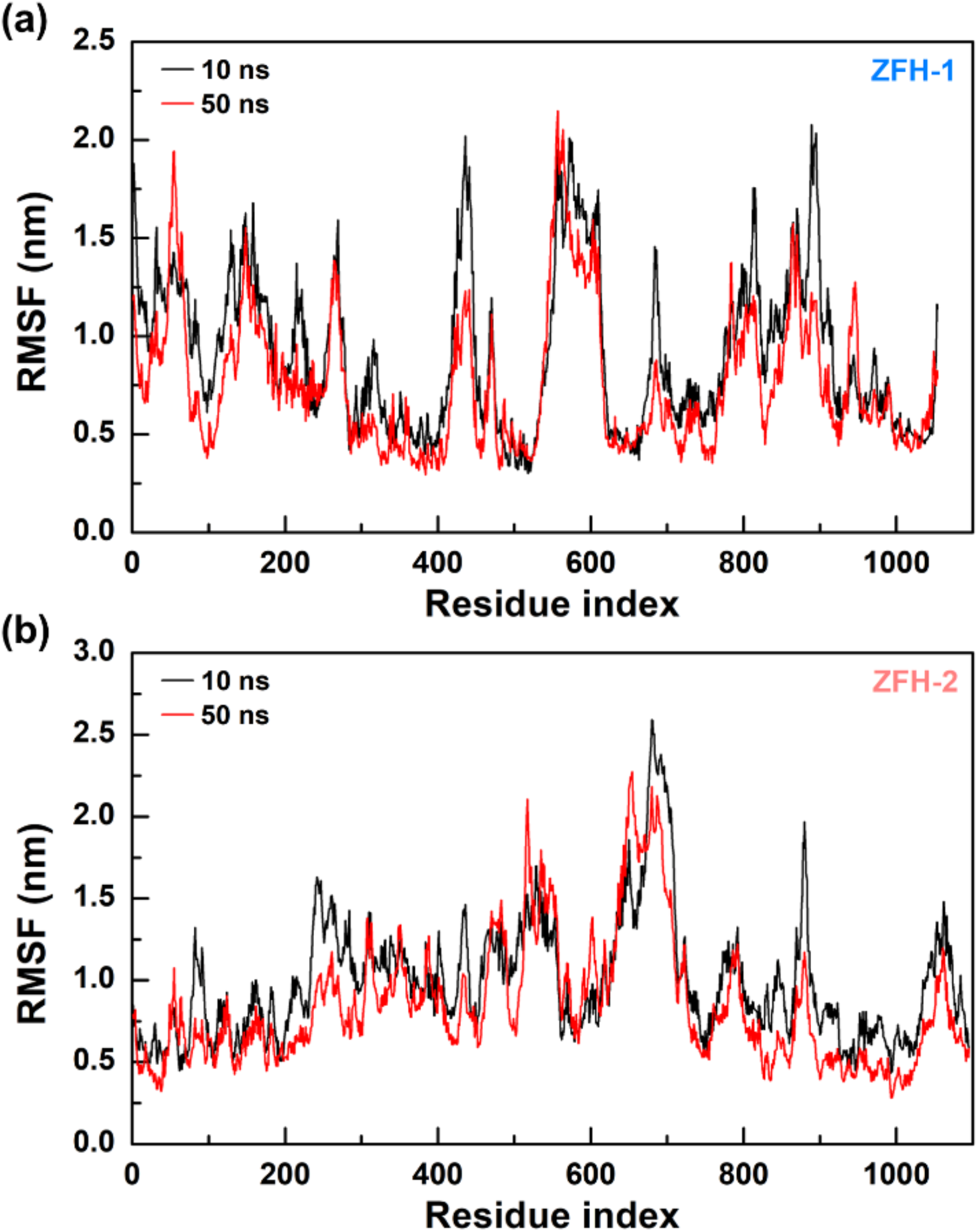
The transmission of the C2H2 zinc figure family transcription factors were instented by the transducted curves for the two transcription factors as ZFH1 in blue and ZFH2 in red.

**Figure 7.**
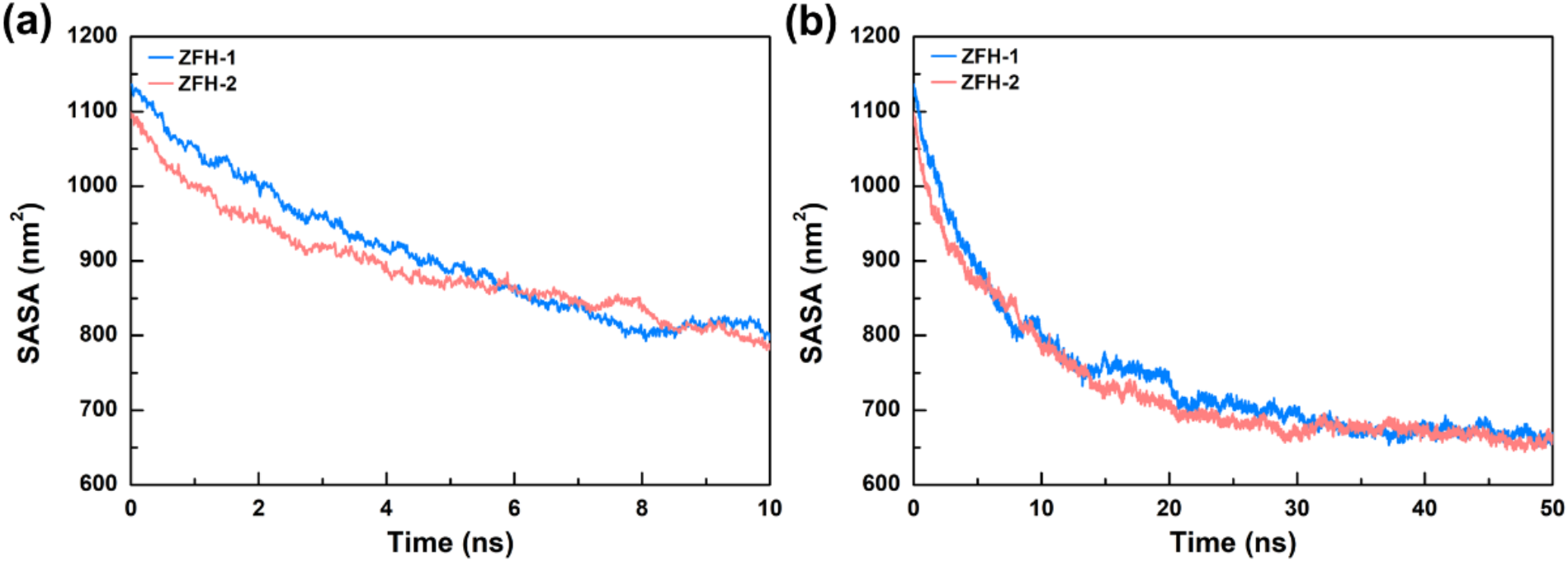
The transmittive transductive conformative transdudite curves for interreaction of the transcription factors as in different curve standing for ZFH2 in comparison with ZFH1 in red and ZFH1 in blue.

Comparably, the conformational exchange were less frequency of the two TFs of the zinc finger family with similar increasing trend. Particularly, most obvious phenomena between the two type of transcription factors were constrained in the RMSF particularly in the DNA binding motif with nearly one magnitude increase in the most highly freedom region as that in the C-terminal of Rex and that in the DNA binding region or the loop region of the two TFs of zinc finger family, respectively, suggesting potentially different molecular mechanism that involved in the DNA recognization and induced conformational change exerted to the DNA by the TFs. The most distinguish property between the two TFs of the C2H2 zinc family with the Rex regulator reflected by the dynamic simulation was solvent accessibility surface area that indicate the almost constant volume exchange of the Rex but sharply decreased of the volume for ZFH1 and ZFH2 which were approached stable until 30ns upon the thermol impact. In conclusion, the above results revealed the compact and stable structure of two TFs of the zinc finger family may available for the functional performance, moreover, significant differences of the main properties demonstrated variety of regulation mechanisms of different elements induced by the TFs with different DNA binding motifs.

## Discussion

The transcription regulation is the most frequently investigated scientific issues that date back to several centuries ago yet the molecular mechanisms that related to the fundamental principles and the detailed recognition properties that remain under completely and precisely understanding since the complicated regulatory system and the lack of efficient research techniques during a long period of time. The main purpose of this article was to demonstrate the entire structural movement of the typical transaction factors that participate in the different biological pathways in regulating speculated elements in the genome of the modeling species as a short cut review for the insight into the propounding impact of the thermal dynamic on the structural remodeling of the protein and DNA, suggesting the potential instructions of the proteins and the particular residues that were included in the binding site on the transcription initiation of the target gene from the specific regulation elements. The research of this article was essentially based on the molecular dynamic simulation method that elucidated the potential characteristics of the entire protein especially the particular domains that were typically DNA binding motifs in the recognition and association of the TFs with the experimentally determined or proposed DNA fragments, inferring the magnetic consequences of proteins on the changeable progression of the elements induced by the atmosphere interactions between the animo acid residues and the nucleotides. These included the following results in details. Firstly the initial conformation of the transcription factor that was belonged to the redox sensing regulator with the helix loop helix motif that were universally presented in various kinds of TFs were assigned for the dynamic simulation. This was transformed into the root mean square deviation (RMSD) to reflect the overall conformational change of the protein, DNA and the comparison of the stability upon DNA association including the simulation of 10ns and 50ns, suggesting the inflection of the protein structure by the modulated DNA fragments as well as the double helix opening induced by the protein in particular by the interacting amino acid residues that are dynamically regulated the structure of the DNA, which was also indicated by the RMSF values in which the highly freedom of bases were just located in the modulated DNA binding site that was not constantly but flexibly consisted by variable residues and nucleotides except for some of the stable ones. The solvent accessibility surface area (SASA) was in consistent with the above phenomenon in conclusion with the stabilization of the TF by the DNA association. Further analysis of the interacting pairs of the DNA associated complex also confidentially reveals the identity of the individual residues and nucleotides that interact with each other in a illustrated form and the RMSD variation per each of the residues of the protein that contributed most to the formation of the complex while with different dynamic properties in consistent with the above analysis and investigated the molecular mechanisms regulating to the DNA remodeling in the elements and orientating the open of the double helix that may deduce stepwise furthermore conformational changes to hinder or activated the gene transcription through secondary or even tertiary structure reassembly of the DNA sequence. Although the increased stability and the reduced freedom of the protein especially for the amino acid residues in the binding site due to the majority of the stable interactions between the residues and the nucleotides but unstabilized the interactions between the base pairs, resulting the conformational change of the DNA binding site and the relative positional change of some of the protein residues involved in the interaction with the bundle of the interacted single strand of the DNA, which probably inferred the molecular mechanisms that the continuous and enlarged conformational change finally leading to the release of the transcription factors from the DNA strands, suggesting the fundamental rule of the DNA recognition and deassociation.

## Method

The results in this article were assigned by the following protein modeling methods including the prediction of the protein structure in the algorithm designed by the scientists from the collaborated company as well as the protein Docking method described in the former articles and developed by the scientists from the above company. The protein simulation was performed by the specialized team of the above company in collaboration with the researchers in our lab through the different types of softwares installed with various simulation packages, for example the Gromacs algorithms. The protein simulation method was initially established by the researchers from the specialized group through the Monte Carlo algorithms and developed by researchers from various groups including those from ^9^and^10^ to improve the efficiency and performance of the simulation systems. This was now achieved by the Gromacs algorithms and so forth.

The dynamic simulation in this article was performed by introducing the developed Gromacs program^10^ under the constant pressure and temperature based on the Amber14SB^11^ force field with TIP3P^12^ solvent model. In the dynamic simulation process the constrain^13^ of the hydrogen bonds was performed with Gromacs algorithms in the time step of 2fs, the other parameters^14-16^ used were listed in the supplementary data, Table S1. The systems mentioned in this article were assigned for the dynamic simulation in the time interval of 50ns, the simulation results were saved every 10ps to present the configuration of the systems at different time points with the equilibration time of 1ns prior to the following simulation steps. The data set was output by the specialized programs installed in the Gromacs and the VMD software for the display of the simulation results. The RMSD, RMSF, SASA and other properties that are used for the representation of the simulation data flow were also output by the above software and so forth. The protein modeling were performed with Alphafold^17^.

## Data availability

Thanks for the supporting from the research group of ^(18)^ that providing and allowing the usage of the relative structures^(18)^ and for the publicative reuse in this article. Thanks for the methodology refinement of the supporting from ^(8)^

## Contribution

Dr. F. Jin designed and manipulated the project including the creation of research spirits, design of the experimental and simulation protocols, as well as the analysis and the evaluation of the results and part of the data flows. The specialized group from the company of Phadcalc manipulated the protocols of protein modeling, dynamic simulation, data analysis and the display of results. The researchers in our lab participated in the experimental study, part of the data analysis and the results display. The entire project was conducted and supervised by the team leader of our lab, Professor K. Xu.

## Acknowledgment

Thanks for the research group from the^18^ that provided and allowed the use of the structures that were published before and deposited in the protein databank. Special thanks for the research group of our lab in collaboration with that in the Phadcalc company. Thanks for the assistance of the specialists in the collaborated groups in particular the research scientist Mr Li and the specialized researchers in the groups that participate in the protein modeling method development as well as the protein docking in subsequently development of the Gromacs based molecular simulation. This project was also assisted by the Supercomputer research groups in collaboration with the researchers particular Miss Liu in the collaboration groups. In addition this group were also assisted by the supercomputer in the research group in collaboration with the work stations in our lab as well as in the compony for special thanks. In particular the specialists from our university (Tongji University) were many thankful.

## Founding

This work was funded by grants from the National Key Research and Development Program of China (2021YFA1302200 to K. X.), Shanghai Sailing Program (20YF1453100), Shanghai Health Commission Clinical Research Special Youth Project (20204Y0320), the Thousand Talents Plan-Youth to K. X., and Fundamental Research Funds for the Central Universities (Tongji University).

