## Supplemental figure and tables for "Structural and modelling insights into the dynamic association between the transcription factor and DNA"

Supplementary data

Figure S1. The predicted structure of the transcription factors of the C2H2 zinc finger family, separately in the cartoon form through protein modeling algorithm. .

Supplementary tables

Table S1 The simulation parameters included in the simulation process were listed below

Table S2 The predicted DNA sequence that was potentially recognized by the transcription factor of C2H2 zinc finger family

.

Supplementary figure

Figure S1

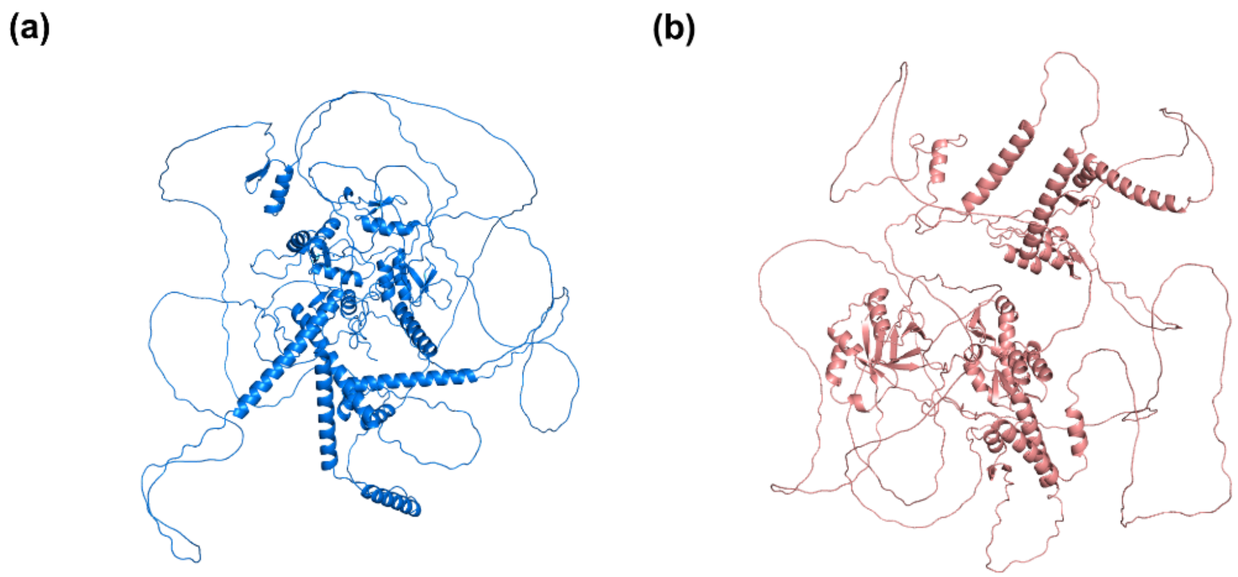


Supplementary table

Table S1

| Parameters | Values |
|---|---|
| Simulation time | 10ns, 50ns |
| Step | 2fs |
| Electrostatic interactions | Particle–mesh Ewald |
| Contact distance | 10ai |
| Minimization | Steepest descent |
| Temperature | 298.15K |
| Pressure | 1bar |
| Algorithm | NVT, NVP |
| Equilibration time | 1ns |

Table S2

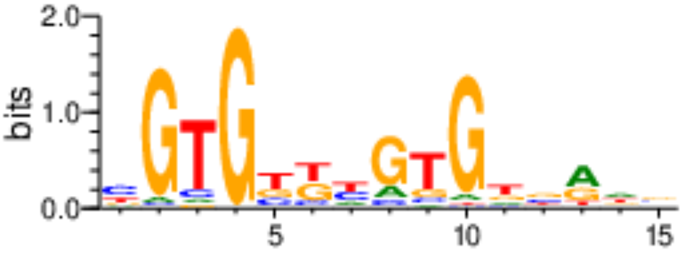
2.0
1.0
0.0
bits
5
10
15

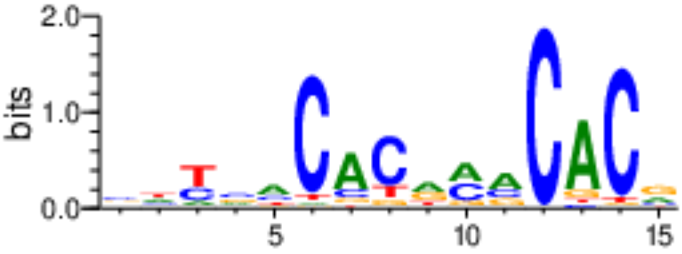
2.0
1.0
0.0
bits
5
10
15
